# Multiple linear regression allows weighted burden analysis of rare coding variants in an ethnically heterogeneous population

**DOI:** 10.1101/2020.06.11.145938

**Authors:** David Curtis

## Abstract

Weighted burden analysis has been used in exome-sequenced case-control studies to identify genes in which there is an excess of rare and/or functional variants associated with phenotype. Implementation in a ridge regression framework allows simultaneous analysis of all variants along with relevant covariates such as population principal components. In order to apply the approach to a quantitative phenotype, a weighted burden score is derived for each subject and included in a linear regression analysis. The weighting scheme is adjusted in order to apply differential weights to rare and very rare variants and a score is derived based on both the frequency and predicted effect of each variant. When applied to an ethnically heterogeneous dataset consisting of 49,790 exome-sequenced UK Biobank subjects and using BMI as the phenotype the method produces a very inflated test statistic. However this is almost completely corrected by including 20 population principal components as covariates. When this is done the top 30 genes include a few which are quite plausibly associated with the phenotype, including *LYPLAL1* and *NSDHL*. This approach offers a way to carry out gene-based analyses of rare variants identified by exome sequencing in heterogeneous datasets without requiring that data from ethnic minority subjects be discarded. This research has been conducted using the UK Biobank Resource.

## Introduction

We have previously developed a method of weighted burden analysis which allows all variants within a gene to be included in a case-control analysis to test whether there is on average an excess of highly weighted variants among cases. In the original conception, implemented in the SCOREASSOC program, both common and rare variants were included but a parabolic function based on minor allele frequency (MAF)was applied such that rare variants would be assigned a higher weight than common ones (Curtis, 2012). This approach was subsequently extended in a number of ways. Functional weights were assigned based on the predicted effect of each variant on the function of the gene, so that for example variants predicted to produce a truncated protein product would be weighted more highly than synonymous variants, and an overall weight for each variant was derived as the product of the frequency and functional weights (Curtis, 2016). Unlike other approaches applied to exome sequence data, this means that all variants can be included in a single combined analysis without having to dichotomise according to allele frequency or predicted impact (Cirulli et al., 2020; Zhao et al., 2020). Sets of genes within a metabolic pathway could be jointly analysed by testing whether the overall burden of rare, functional variants varied between cases and controls across the set of genes rather than within an individual gene (Curtis, 2016). The comparison of variant burden was then implemented in a ridge logistic regression framework with and this allowed the inclusion of covariates such as population principal components as well as additional risk factors such as pathogenic copy number variants and polygenic risk scores (Curtis, 2019). These approaches were applied to large samples of exome sequenced cases and controls and implicated genes affecting functioning of the glutamatergic NMDA receptor in schizophrenia and genes coding for tyrosine phosphatases in late onset Alzheimer’s disease (Curtis et al., 2019, 2018).

Additional exome sequenced datasets are becoming available and some of these, such as the UK Biobank, have been phenotyped for a number of quantitative traits (Hout et al., 2019). It is plausible that variants disrupting the functioning of particular genes might be associated with changes in the mean value of some of these traits and an obvious approach would be to carry out linear regression rather than logistic regression in order to test for this. However there are some important considerations which need to be addressed.

In contrast to a targeted case-control study, biobank samples may not be ethnically well-matched and this is expected to impact testing for an excess of rare, functional variants in exome-sequenced sample. Even in studies which seek to match cases and controls there may be residual stratifications which affect the results and this occurred in the Swedish schizophrenia study (Genovese et al., 2016). Here, a higher proportion of cases than controls had a substantial Finnish component to ancestry and at the same time there was a lower frequency of rare, damaging variants in those with Finnish ancestry generally, across both cases and controls. In fact, there was an excess of rare damaging variants among the schizophrenia cases but this only became apparent when ancestry was included as a covariate or when the analysis was restricted to subjects without Finnish ancestry (Curtis et al., 2018; Genovese et al., 2016). This provides a practical example of how unrecognised population stratification may distort the results of rare variant burden analyses. Of particular relevance to biobanks is that there is an expectation that such stratification will produce artefactual results even if there is in fact no difference in the general distribution of variant allele frequencies between subjects with different ancestries. If the bulk of subjects share similar ancestry but there is a minority cohort with different ancestry then we can expect by chance that some variants will have different frequencies between the main and minority cohorts. However, a variant which is relatively common in the minority cohort will have a lower frequency in the sample overall. This results in the expectation that there will appear to be an excess of apparently rarer variants in minority cohorts. If the phenotype in question has a different mean value in the minority cohort then there will be an artefactual association between the phenotype and rare variants, whether rarity is defined using a threshold or is used in a weighting scheme.

One approach to dealing with heterogenous ancestry within biobank samples is to restrict attention only to those of a particular ethnicity. The UK Biobank sample contains 503,317 subjects of whom 94.6% are of white ethnicity, somewhat higher than for the population as a whole (Fry et al., 2017). Regrettably, it seems to have become standard practice to simply ignore the non-white subjects. To give two recent examples, a genome-wide meta-analysis of problematic alcohol use only considered 435,563 European-ancestry individuals and a genome-wide association study of susceptibility to keratitis only considered 337,199 subjects of European ancestry (Xu et al., 2020; Zhou et al., 2020). Likewise, a recent analysis of the 49,960 exome-sequenced UK Biobank subjects only used information from 45,596 European subjects (Zhao et al., 2020). Restricting attention to subjects with white European ancestry within UK Biobank discards information from thousands of citizens who have volunteered personal information, donated biological samples and undergone uncomfortable investigations with the aim of contributing to knowledge about disease. We believe that simply ignoring data from ethnic minority subjects is ethically indefensible (Curtis and Balloux, 2020).

A second issue to be addressed in weighted burden testing is the nature of the weighting which is to be applied according to MAF. In its original conception, the method was intended to incorporate variants of all allele frequencies and an example application used Crohn’s diseases as the phenotype, since susceptibility is influenced by both common and rare variants (Curtis, 2012). However over the course of countless genome wide association studies it has become apparent that it is in fact very unusual for common variants to exert substantial effects on risk and when dealing with next generation sequence data it does not make sense to include common variants since they will essentially produce noise which may swamp any real signal. However with the weighting scheme originally proposed, described by a parabola with value of 1 at MAF = 0.5 increasing to 10 at MAF ~= 0, the allocated weight is almost the same for a variant with MAF = 0.01 as it is for ultra-rare variants. Since selection pressures mean that very rare variants can have larger effect sizes than less rare ones, it would be desirable to have a revised weighting scheme which would distinguish rare from ultra-rare variants (Long et al., 2017).

A third issue to address when dealing with quantitative traits is how information from different genes within a set should be combined. The original assumptions were that rare variants with a functional impact were more likely to impair normal function of a gene than enhance it and hence that such variants were more likely to be associated with a disease phenotype across different genes. This would mean that variant scores could simply be added across genes within a set (Curtis, 2016). With a quantitative trait we may still expect that rare, functionally impactful variants may impair gene functioning but we need to recognise that impairing the function of some genes may tend to reduce the value of a trait whereas impairing the function of other genes in the same pathway may increase the value. Since the effects may be in opposite directions, we will not wish to simply add up variant scores across related genes unless we can be very confident of the direction of the effect we are expecting. Since we are intending to use this approach for gene discovery we will not generally be in a position to predict the likely direction of effect and so we require a method which is agnostic regarding this.

Here we develop the weighted burden approach previously applied to case-control data in three different ways. We produce a modified scheme for weighting by allele frequency to give additional weight to extremely rare variants. We implement linear regression rather than logistic regression and incorporate population principal components as covariates. We combine evidence across different genes within a set using p values rather than variant scores. We apply the revised method to analyse a quantitative trait, BMI, in an ethnically heterogeneous dataset, the UK Biobank.

## Methods

### Variant weight according to allele frequency

The original formula we proposed for assigning a weight according to allele frequency was

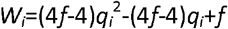

where *q_i_* is the allele frequency of the *i*th variant and *f* is a weighting factor equal to or greater than 1. The formula describes a parabola with a minimum value of 1 at *q*=0.5 and rising to *f* at *q*=0 and *q*=1. In typical applications a value of 10 would be chosen for *f*.

In order to provide a scheme which more strongly distinguishes rare from very rare variants, we will choose *q*’ to mean the frequency of the rarer allele and we will set a threshold of *t* for the MAF. We will either discard variants with *q*’>*t* of else we will assign them a weight equal to 1. For those with *q*’<=*t* we will assign a weight as:

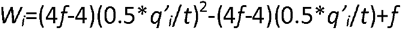

This simply produces weights which increase parabolically from a value of 1 at *q*=*t* or *q*=1-*t* towards a value of *f* at *q*=0 and *q*=1.

### Implementation of linear regression

As described previously, testing for an effect of gene variants on a case-control phenotype involved constructing a weighted burden score for each subject and then carrying out a likelihood ratio test using ridge regression comparing the likelihoods for models which did and did not include the burden scores (Curtis, 2019). Here, we simply calculate the likelihoods using linear regression rather than logistic regression. Each variant is assigned a weight based on its predicted likely effect on gene function and then this functional weight is multiplied by the weight based on the variant frequency as described above. For each subject a gene-wise weighted burden score is derived as the sum of the variant-wise weights, each multiplied by the number of alleles of the variant which the given subject possesses. If a subject is not genotyped for a variant then they are assigned the subject-wise average score for that variant. A number of covariates can be included for each subject, typically consisting of population principal components though if desired factors such as age, polygenic risk score and presence or absence of other known risk factors can also be used. The score and covariates are entered into a standard linear regression model with quantitative phenotype as the outcome variable and after variable normalisation the likelihood of the model is maximised using the L-BFGS quasi-newton method, implemented using the dlib library (King, 2009). The contribution of different variables to risk is assessed using standard likelihood ratio tests by comparing twice the difference in maximised log likelihoods between models with and without the variables of interest. This likelihood ratio statistic is then taken as a chi-squared statistic with degrees of freedom equal to the difference between models in number of variables fitted. The coefficients for each variable can be varied to maximise the likelihood or can be fixed, for example if the effect size of a particular risk factor is known from epidemiological studies. This approach was implemented in a modified version of SCOREASSOC, which outputs the coefficients for the fitted models along with their estimated standard errors and the results of the likelihood ratio test. When association with the gene-wise weighted burden score alone is tested, i.e. when the two models differ only in whether or not the score is included, then the statistical significance is summarised as a signed log p value (SLP) which is the log base 10 of the p value given a positive sign if the score correlates positively with the quantitative phenotype and negative if it correlates negatively. For other analyses the minus log base 10 of the p value (MLP) is output. The support program for SCOREASSOC, called GENEVARASSOC, was also modified to deal with quantitative phenotypes and the linear regression tests.

### Gene set analysis

In case-control analyses the assumption was made that rare, functional variants tended to impair gene function and that impaired gene function increased disease risk, meaning that weighted burden scores could simply be added across genes. For a quantitative trait, impaired function of a gene within a metabolic pathway might either increase or decrease the value of the trait and since genes might have opposite effects it is not appropriate to simply sum the scores of genes within a set. Instead, Fisher’s method for combining p values can be applied. This assumes that, under the null hypothesis that no genes within a set influence the value of the trait, the sum of the natural logs of their *p* values multiplied by −2 will follow a chi-squared distribution with degrees of freedom equal to the twice number of genes. This can conveniently be tested by summing the absolute values of the SLPs and multiplying by −2ln(10) to use as the chi-squared statistic. The statistical significance of the test that one or more genes in the set influence phenotype can then be expressed as minus log base 10 of the p value (MLP) of the chi-squared test.

### Practical application to an example dataset

The UK Biobank dataset was downloaded along with the variant call files for 49,953 subjects who had undergone exome-sequencing and genotyped using the GRCh38 assembly with coverage 20X at 94.6% of sites on average (Hout et al., 2019). Informed consent from the subjects and ethical approval for research uses of the data had been obtained by UK Biobank. All variants were annotated using VEP, PolyPhen and SIFT (Adzhubei et al., 2013; Kumar et al., 2009; McLaren et al., 2016). To obtain population principal components reflecting ancestry, version 1.90beta of *plink* (https://www.cog-genomics.org/plink2) was run on these variants with the options --*maf 0.1 --pca header tabs --make-rel* (Chang et al., 2015; Purcell et al., 2007, 2009).

The example quantitative phenotype chosen was BMI, which is known to be moderately heritable and which was available for 49,790 subjects. In order to better understand the structure of the data a number of other variables were studied including self-declared ethnicity, birth coordinates within the UK, year of birth and Townsend index of deprivation. The relationships between these variables and the principal components were investigated using multivariate analyses implemented in R and visualised using *ggplot2* and *ggpubr* (https://cran.r-project.org/web/packages/ggpubr/index.html) (R Core Team, 2014; Wickham, 2016).

Analysis was restricted to variants have MAF<=0.01 and weighting based on frequency was applied as described above with a weighting factor, *f*, of 10 and a threshold, *t*, of 0.01. Variants were also weighted according to their functional annotation using the default weights provided with the GENEVARASSOC program, which was used to generate input files for weighted burden analysis by SCOREASSOC. For example, a weight of 5 was assigned for a synonymous variant, 10 for a non-synonymous variant and 20 for a stop gained variant. Additionally, 10 was added to the weight if the PolyPhen annotation was possibly or probably damaging and also if the SIFT annotation was deleterious, meaning that a non-synonymous variant annotated as both damaging and deleterious would be assigned a weight of 30. The full set of weights is shown in Supplementary Table 1, copied from the previous reports which used this method (Curtis et al., 2019, 2018). Variants were excluded if there were more than 10% of genotypes missing in the controls or if the heterozygote count was smaller than both homozygote counts. Each variant was then assigned an overall rate consisting of the product of the frequency weight and the functional weight. For each subject a gene-wise weighted burden score was derived as the sum of the variant-wise weights, each multiplied by the number of alleles of the variant which the given subject possessed.

Gene-wise weighted burden tests for association with BMI were carried out for every autosomal and X chromosome gene listed in the RefSeq GRCh38 release. For each gene, two analyses were carried out. In the first, weighted burden score was used to predict BMI in a simple linear regression model and in the second analysis the first 20 principal components were additionally included as covariates, with likelihood ratio tests used to assess statistical significance as described above.

Gene set analyses were carried out using the 1454 “all GO gene sets, gene symbols” pathways as listed in the file *c5.all.v5.0.symbols.gmt* downloaded from the Molecular Signatures Database at http://www.broadinstitute.org/gsea/msigdb/collections.jsp (Subramanian et al., 2005). For each set of genes the SLPs from the analyses incorporating principal components were combined using Fisher’s method as described above, yielding an MLP as a test of association of the set with the BMI.

## Results

Some principal components were associated with self-declared ethnicity, with the first principal component distinguishing African from European ancestry, the second Asian from European and the fourth South Asian from East Asian (Supplementary Figures 1 and 2). Principal components also varied with place of birth, year of birth and Townsend deprivation index (Supplementary Figures 3 to 6). Likewise, BMI also varied with self-declared ethnicity as shown in Figure 1 and Table 1, as well as with demographic variables (Supplementary Figure 7). Finally, BMI varied with the principal components (Supplementary Figure 8). Thus BMI is a phenotype which can vary between subjects with different ancestries and could be liable to produce artefactual results according to mechanisms such as those outlined above.

**Figure 1.**
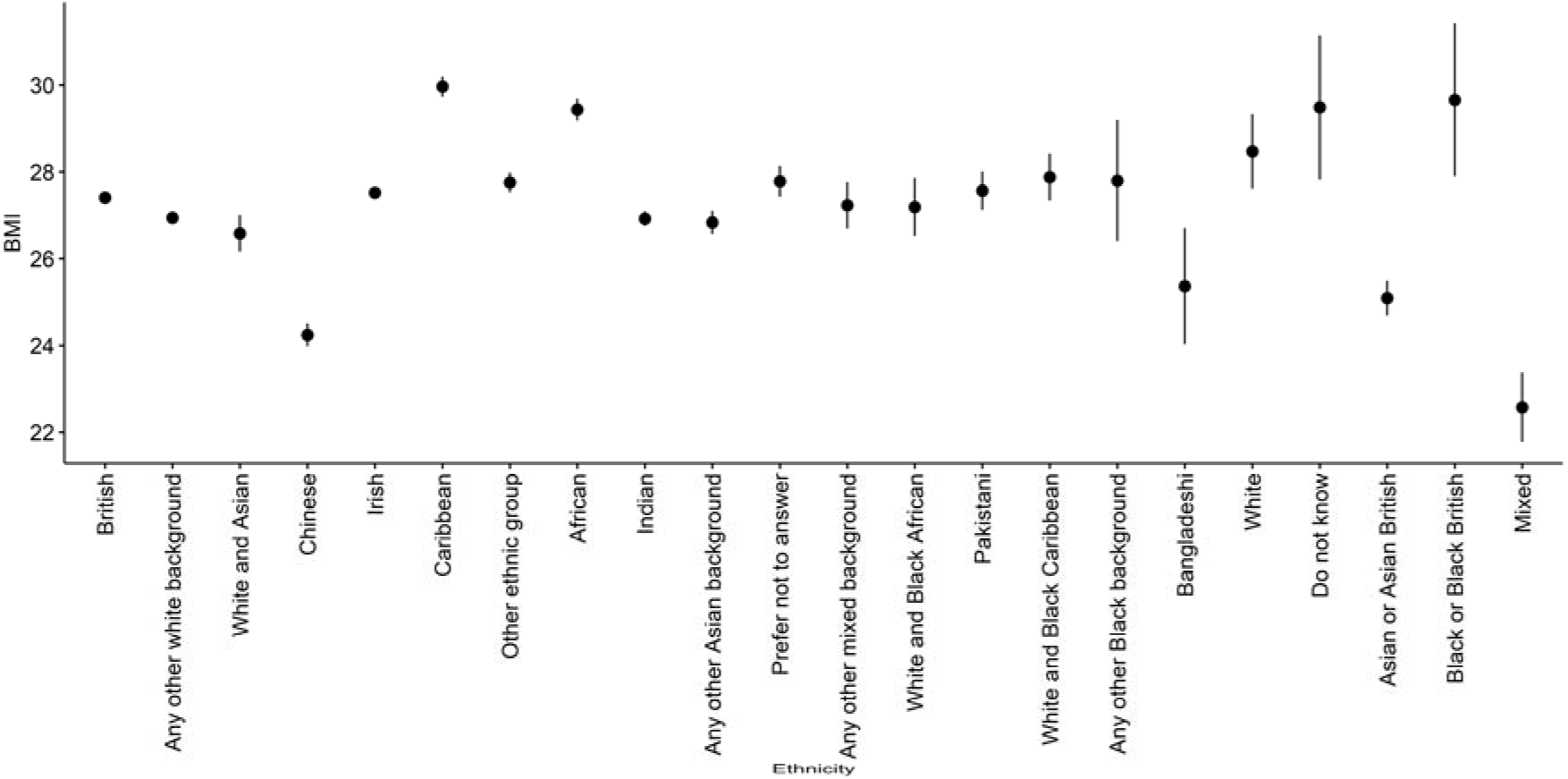
Plot of mean and SE(mean) of BMI against self-declared ethnicity in exome-sequenced UK Biobank subjects.

**Table 1.** BMI versus self-declared ethnicity.

| **Ethnicity** | **Number of subjects** | **Mean (SE)** |
| --- | --- | --- |
| White | 38 | 28.47 (0.86) |
| British | 43254 | 27.40 (0.02) |
| White and Black Caribbean | 96 | 27.88 (0.54) |
| Indian | 696 | 26.92 (0.17) |
| Caribbean | 657 | 29.96 (0.23) |
| Mixed | 2 | 22.57 (0.80) |
| Irish | 1497 | 27.52 (0.12) |
| White and Black African | 48 | 27.19 (0.67) |
| Pakistani | 139 | 27.56 (0.44) |
| African | 337 | 29.43 (0.26) |
| Asian or Asian British | 2 | 25.09 (0.40) |
| Any other white background | 1699 | 26.94 (0.11) |
| White and Asian | 106 | 26.58 (0.42) |
| Bangladeshi | 16 | 25.36 (1.34) |
| Any other Black background | 10 | 27.79 (1.39) |
| Black or Black British | 3 | 29.66 (1.76) |
| Any other mixed background | 118 | 27.23 (0.54) |
| Any other Asian background | 202 | 26.83 (0.27) |
| Chinese | 173 | 24.24 (0.27) |
| Other ethnic group | 502 | 27.75 (0.23) |
| Do not know | 14 | 29.48 (1.66) |
| Prefer not to answer | 174 | 27.78 (0.36) |

There were 21,644 genes for which there were qualifying variants and the QQ plots for the SLPs are displayed in Figure 2. It can be seen that when principal components are not included there is a very marked inflation of the positive SLPs and that this is almost entirely corrected when the principal components are used as covariates. With the principal components included in the analysis, if the highest and lowest 100 SLPs are excluded (since they might capture a true biological effect) then the negative SLPs have an intercept of −0.022 and a gradient of 0.996 while the positive SLPs have an intercept of −0.006 and a gradient of 1.08, indicating only a fairly modest amount of inflation of the positive SLPs.

**Figure 2.**
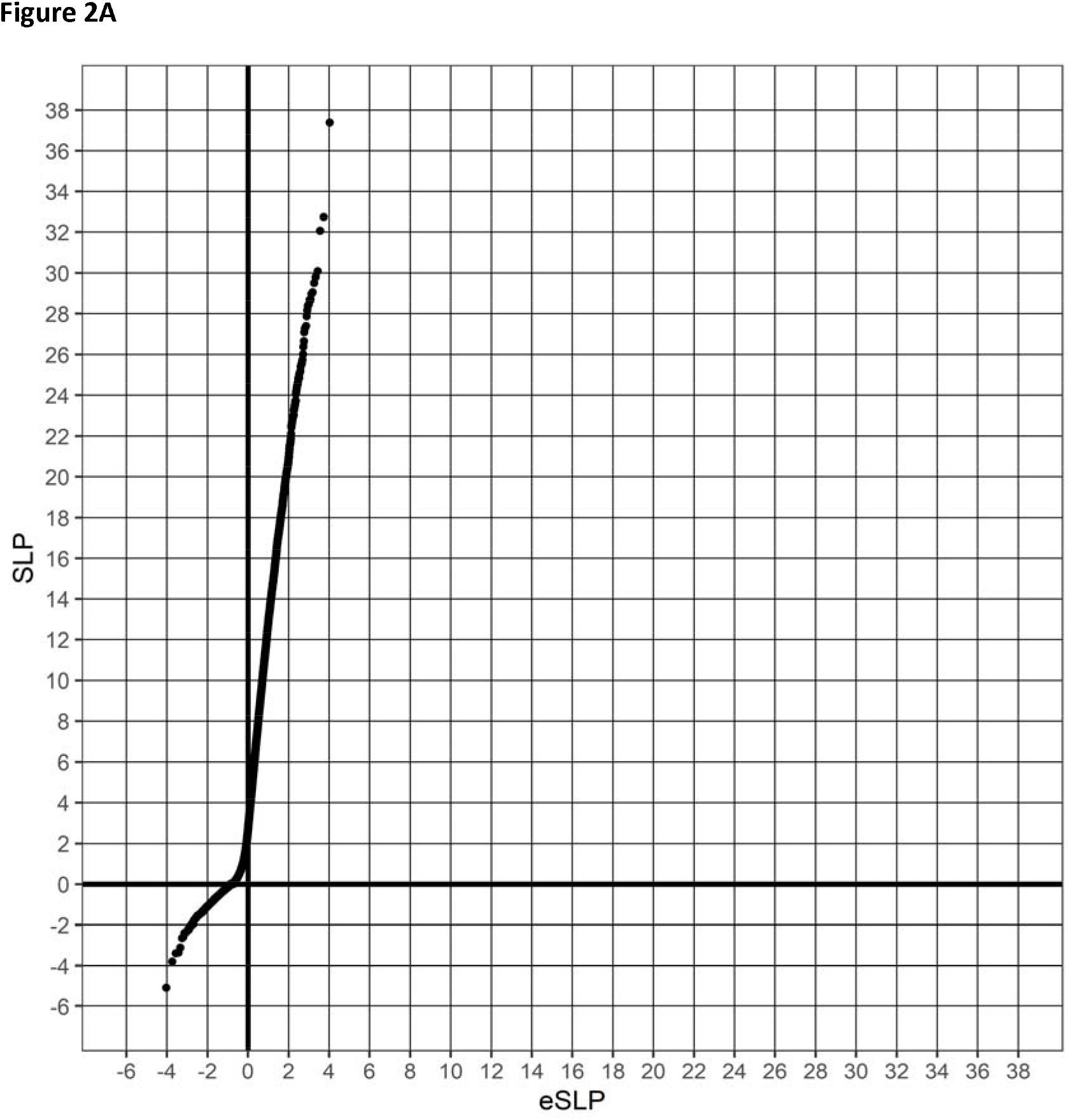
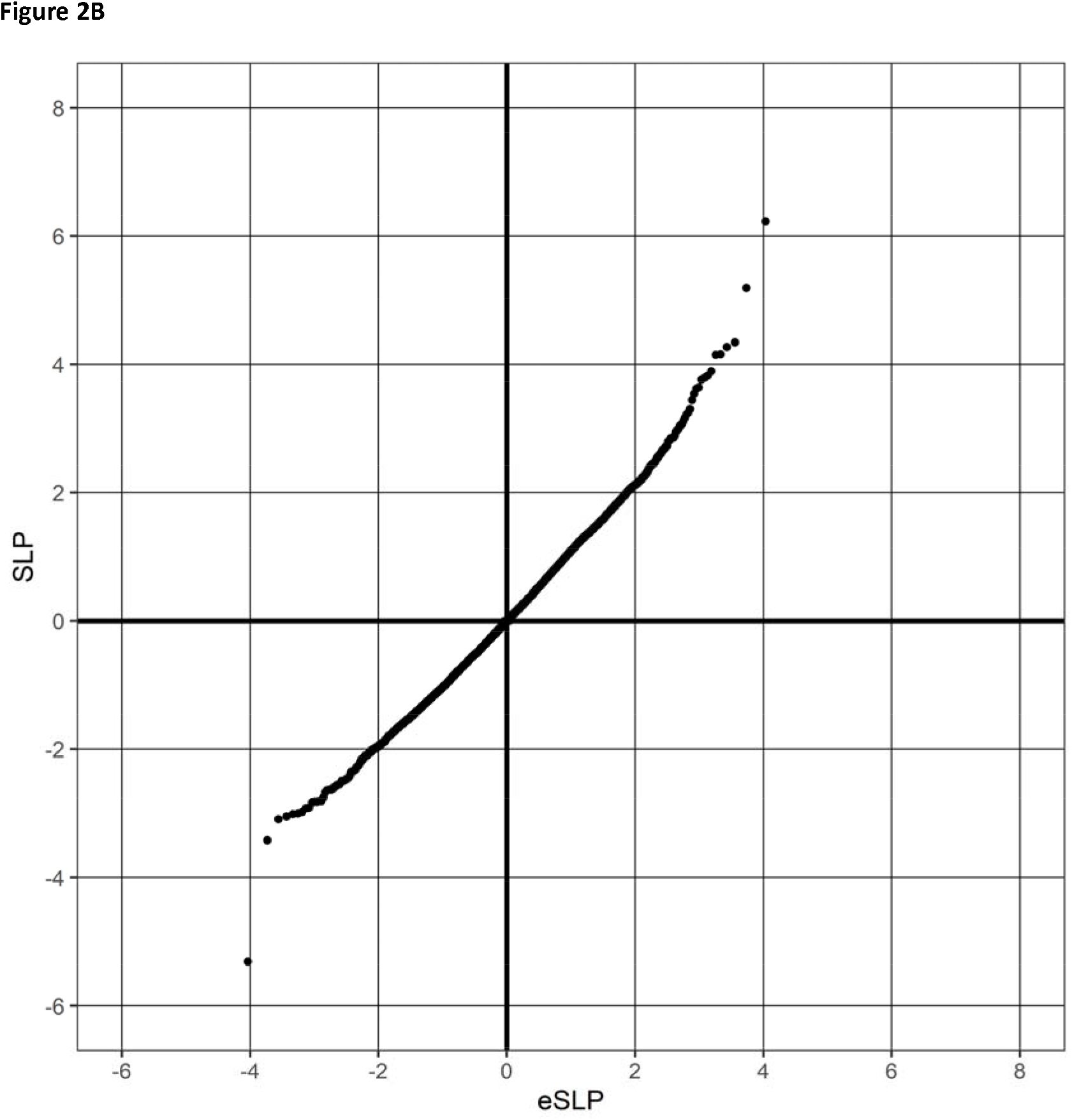
QQ plots of SLPs obtained for weighted burden analysis of 21,644 genes for association with BMI. 1A shows the results for regression of the weighted burden score against BMI alone and 1B shows the results when the population principal components are included as covariates.

Applying a Bonferroni correction for the number of genes tested would mean that the absolute value of the SLP would need to exceed log10(21,644*20)=5.6 or, allowing for the inflation referred to above, log10(21,644*20)*1.08=6.08. This was only achieved by one gene, *CCDC140*, which codes for a small effector of CDC42 and does not seem a particularly plausible candidate to have a direct biological influence on BMI. If the test were well behaved then under the null hypothesis one would expect 11 genes to have SLP of 3 or more and 11 to have SLP of −3 or less. The observed numbers are 22 and 7, possibly reflecting the modest inflation of the positive SLPs. Genes with absolute value of SLP of 3 or more, equivalent to p<=0.001, are listed in Table 2. Although these do not meet formal standards of statistical significance after correction for multiple testing, there are a few for which disruption of functioning might plausibly affect BMI.

**Table 2.**
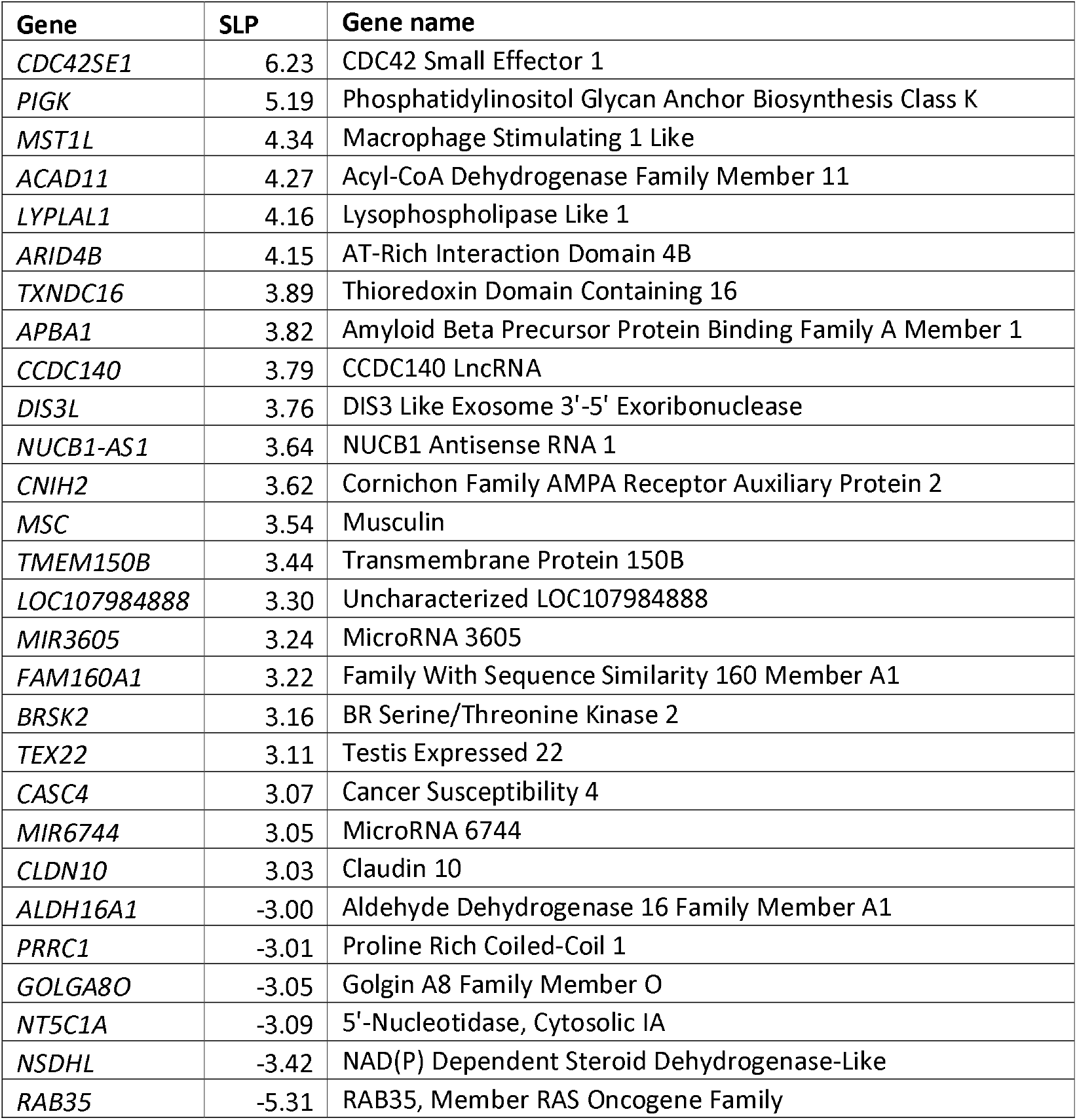
Genes with absolute value of SLP of 3 or more (equivalent to p<=0.001) for test of association of weighted burden score with BMI.

The result for *ACAD11* may be of some interest because its product is involved in mitochondrial beta-oxidation of lipids and energy production. Additionally, two common variants in *ACAD10*, which has a similar function, are associated with obesity and type 2 diabetes in Pima Indians and *Acad10* knockout mice have excess weight gain which increases with age (Bian et al., 2010; Bloom et al., 2018). However in the present study the SLP for *ACAD10* itself is only −0.51.

There is good evidence from previous GWASs that common variants in or near *LYPLAL1* are associated with a variety of metabolic traits including central obesity, fatty liver and waist-to-hip ratio, although ablation of *Lyplal1* in mice does not lead to any significant abnormality in phenotype or metabolic physiology (Watson et al., 2017). *LYPLAL1* is also one of the top 10 genes implicated in insulin resistance from two GWASs (Lotta et al., 2017; Scott et al., 2012). Knock out of *LYPLAL1* has recently been shown to cause reduced insulin-induced AKT2 phosphorylation and glucose uptake in human adipocytes (Chen et al., 2020).

The product of *BRSK2*, SAD-A kinase, is involved in the regulation of pancreatic islet β-cell size and mass and insulin secretion in response to glucose levels (Nie et al., 2013). However there does not seem to be any other evidence that it might have an influence on BMI.

*NT5C1A* codes for a 5□-nucleotidase which catalyses the hydrolysis of AMP and silencing it in mouse tibialis anterior muscle results in reduced protein content and increased glucose uptake (Kulkarni et al., 2011).

The product of *NSDHL* is involved in cholesterol biosynthesis and different allelic variants in it are known to cause the X-linked disorders CK syndrome, characterised by intellectual disability and an asthenic build, and CHILD syndrome, characterised by hemidysplasia, erythroderma and limb defects, which is typically lethal in males (McLarren et al., 2010; Ramphul et al., 2019). A non-synonymous polymorphism in *Nsdhl* has been reported to be associated with reduced HDL cholesterol levels in mice (Bautz et al., 2013).

Among other functions, *RAB35* may regulate the insulin-stimulated translocation of glucose transporter SLC2A4/GLUT4 in adipocytes and SNPs near *RAB35* are associated with backfat thickness at 100 kg in pigs (Chen et al., 2019; Davey et al., 2012).

The results from the gene set analyses, are summarised in Figure 3A. After the top 20 scoring sets are removed the MLPs have a gradient of 1.26 with an intercept at 0.02, indicating quite marked inflation. It seemed possible that this might represent the cumulative effect of the modest inflation noted in the individual SLPs and so the procedure of combining them according to Fisher’s method was repeated after first dividing them by the inflation factor of 1.08. This produced the results displayed in Figure 3B. Here the gradient is 0.70 with an intercept at −0.03, indicating some deflation of the statistic. Applying a Bonferroni correction for the number of sets tested would mean that the value of the MLP would need to exceed log10(1,454*20)=4.5 and this was achieved by only one set, PHOSPHOINOSITIDE_BINDING with MLP=4.71. This set includes 19 genes and the result is driven by the fact that it contains both *PIGK* (SLP=4.81) and *RAB35* (SLP=-4.92). Only two other genes in the set are individually significant at p<0.05, *CYTH3* (SLP=-1.55) and *ZFYVE16* (SLP=-1.38). Although all these genes fall into the category of phosphoinositide binding they do not seem to be involved in any shared metabolic processes and, apart from *RAB35*, their functions are not such that one would expect them to have a direct effect on BMI. There were 8 gene sets with MLP>2 (equivalent to p<0.01) and these are listed in Table 3. None look particularly plausible as candidates to be involved in directly influencing BMI.

**Figure 3.**
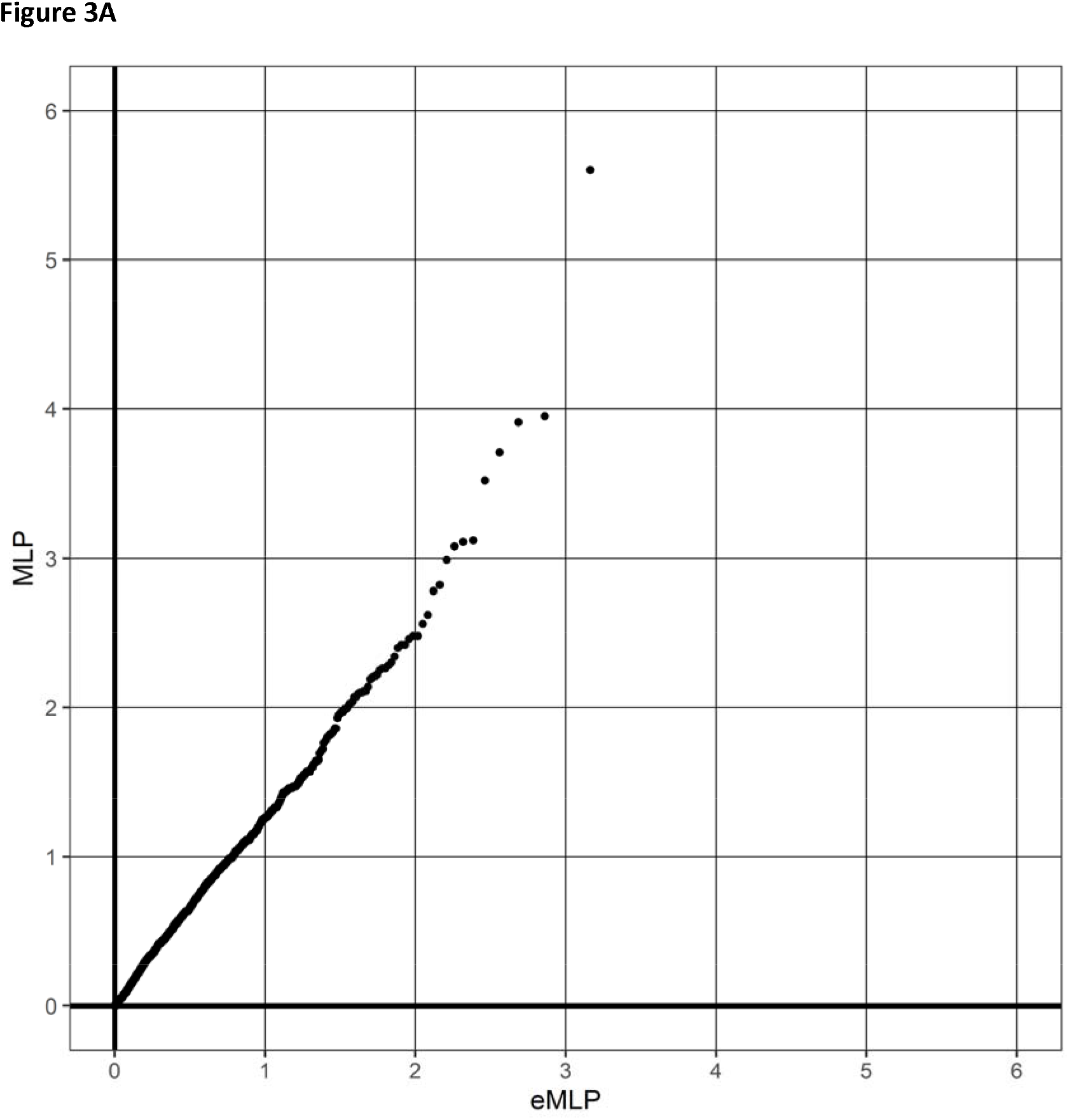
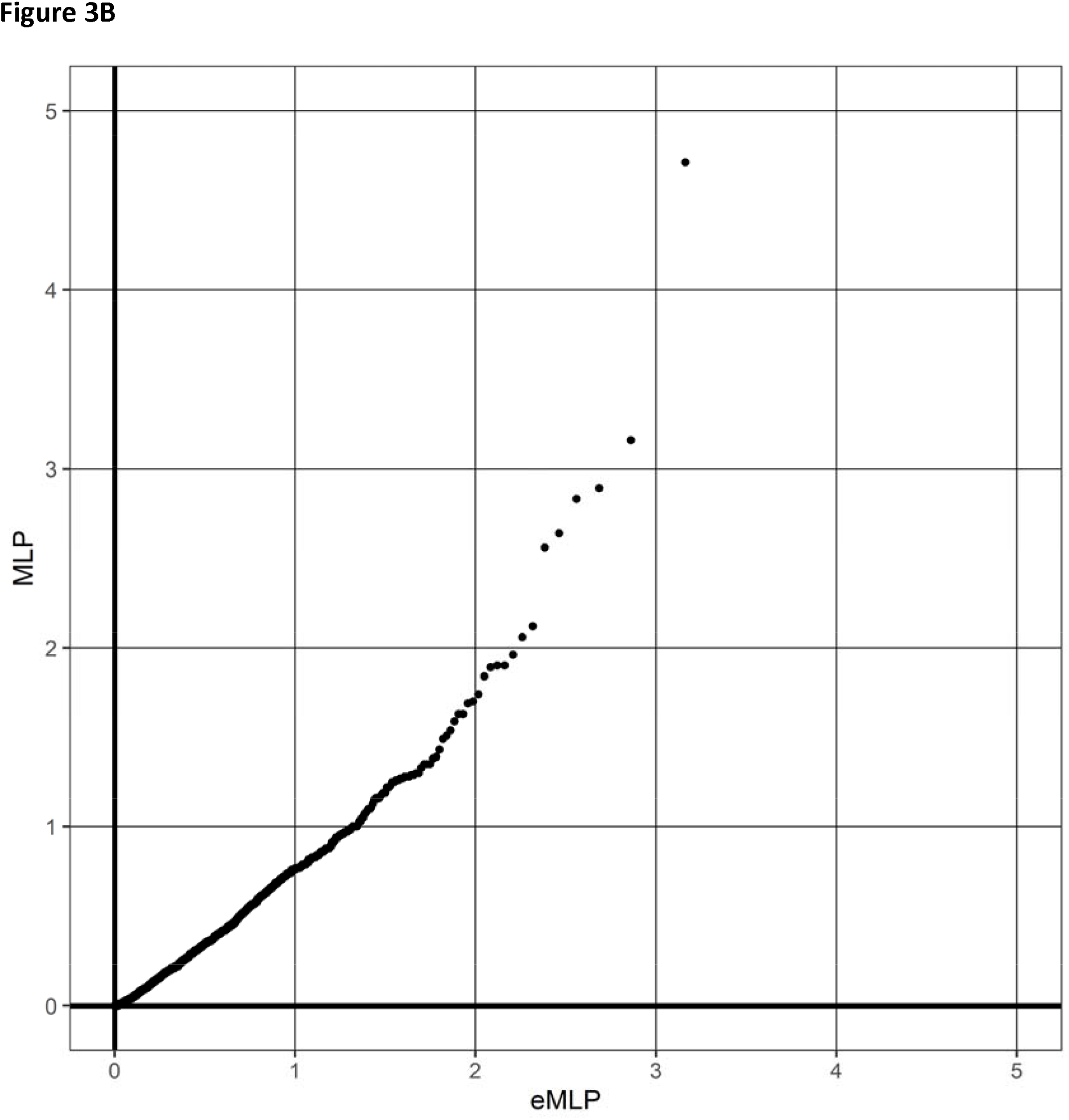
QQ plots of MLPs for 1,454 gene set analyses performed by combining SLPs obtained from the gene-wise analyses incorporating population principal components. 2A shows the results obtained for the uncorrected analyses and 2B shows the results obtained after first dividing the SLPs by an inflation factor of 1.08.

**Table 3.** Gene sets with value of MLP of 3 or more (equivalent to p<=0.01) for test of association of weighted burden scores with BMI.

| Gene set | MLP |
| --- | --- |
| PHOSPHOINOSITIDE_BINDING | 4.71 |
| CYTOKINESIS | 3.16 |
| CELL_DIVISION | 2.89 |
| CLATHRIN_COATED_VESICLE | 2.83 |
| PHOSPHOLIPID_BINDING | 2.64 |
| ENDOCYTIC_VESICLE | 2.56 |
| COATED_VESICLE | 2.12 |
| EXONUCLEASE_ACTIVITY | 2.06 |

## Discussion

The results demonstrate that weighted burden tests are very sensitive to artefactual false positive results arising from population stratification even when, as here, only a small proportion of subjects report a minority ethnicity. Perhaps surprisingly, this problem is almost completely resolved by inclusion of population principle components as covariates. In fact, population stratification is intrinsically less challenging for weighted burden approaches than for variant-wise association analyses because artefacts arise from differences in the distribution of allele frequencies rather than from the differences in frequencies of individual alleles. Hence, inclusion of the principal components as covariates seems to be quite effective. By contrast, principal components are well-recognised not to fully capture population structure in the context of variant-wise tests of association and polygenic risk scores (Lawson et al., 2020). It may be worth noting that in the current dataset some categories of self-reported ethnicity included very small numbers, meaning that it would not be possible to use these categories as covariates. Rather, the principal components seem to adequately capture ethnicity and other sources of stratification.

In terms of the findings from the example analysis, only one gene (just) reaches conventional criteria for genome-wide significance and from what is known of its function it does not seem likely that this represents a real biological effect. On the other hand, there are a few highly ranked, though not formally statistically significant, genes where it is quite plausible that rare, functional variants might be exerting an influence on BMI. Perhaps the two most notable examples would be *LYPLAL1*, for which it is well established that common variants are associated with obesity and related traits, and *NSDHL*, for which it is known that some rare allelic variants can produce a syndrome with asthenia as part of the phenotype. These findings could be explored in other datasets and it is reasonable to expect that as sequence data becomes available for additional UK Biobank subjects it will be possible to distinguish true positive results. The analysis of gene sets has on this occasion failed to yield any further insights or even any suggestive findings. It is possible that such analyses might be more successful when applied to other phenotypes or with gene classifications which better captured biological function.

To conclude, the approach presented here seems to show promise as being statistically fairly well-behaved and, importantly, as allowing heterogeneous datasets to be analysed without having to discard data from ethnic minority subjects.

## Software availability

SCOREASSOC and GENEVARASSOC along with related scripts and documentation can be downloaded from https://github.com/davenomiddlenamecurtis?tab=repositories.

## Data availability

The raw data is available on application to UK Biobank. The summary gene-wise results for all analyses are included in the supplementary file *UKBB.BMI.allSLPs.20200611.txt*. Detailed results with variant counts cannot be made available because they might be used for subject identification.

## Acknowledgments

This research has been conducted using the UK Biobank Resource. The author wishes to acknowledge the staff supporting the High Performance Computing Cluster, Computer Science Department, University College London. This work was carried out in part using resources provided by BBSRC equipment grant BB/R01356X/1. The author declares he has no conflict of interest.

## Supplementary material

**Supplementary Table 1.**
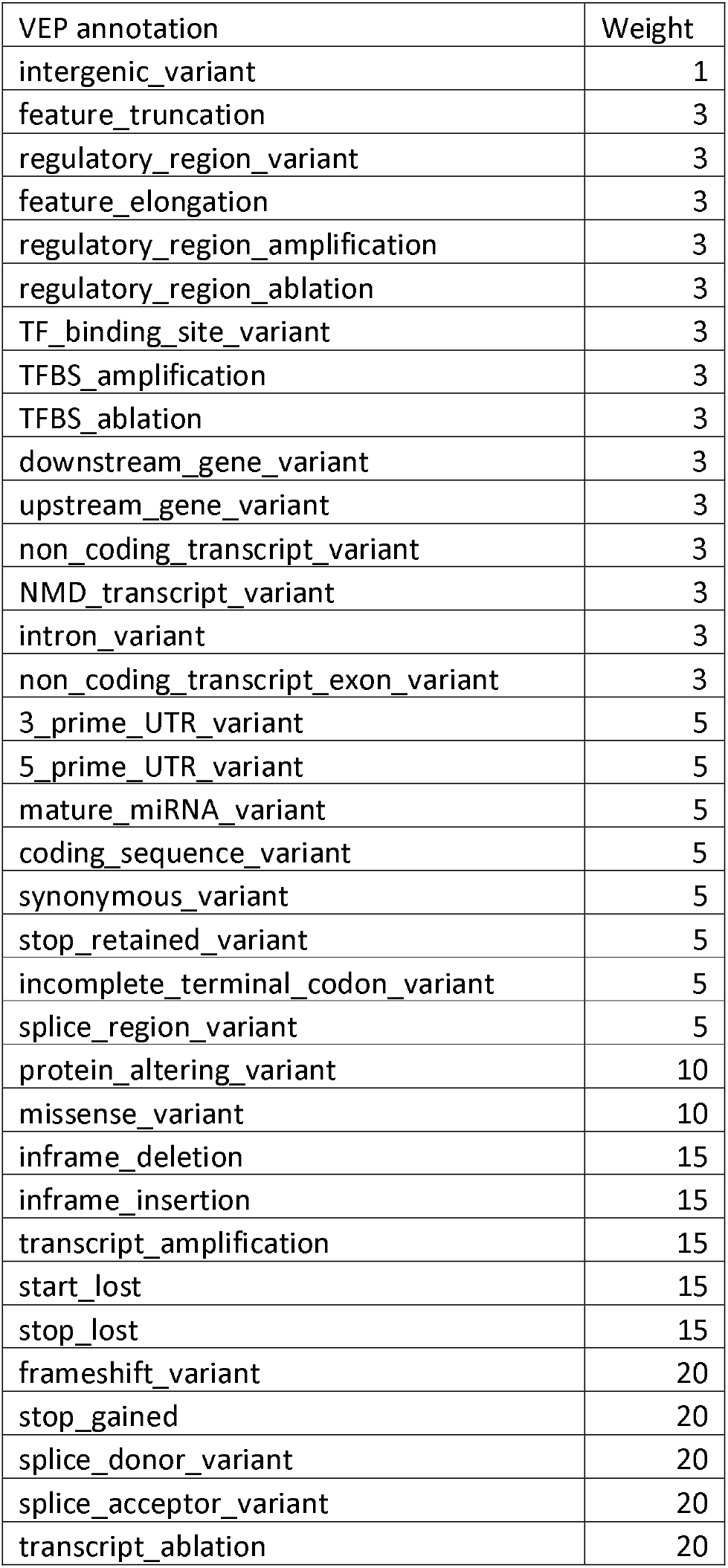
The table shows the weight accorded to each type of variant as annotated by VEP (McLaren et al., 2016). 10 was added to this weight if the variant was annotated by Polyphen as possibly or probably damaging and 10 was added if SIFT annotated it as deleterious (Adzhubei et al., 2013; Kumar et al., 2009).

| VEP annotation | Weight |
| --- | --- |
| intergenic_variant | 1 |
| feature_truncation | 3 |
| regulatory_region_variant | 3 |
| feature_elongation | 3 |
| regulatory_region_amplification | 3 |
| regulatory_region_ablation | 3 |
| TF_binding_site_variant | 3 |
| TFBS_amplification | 3 |
| TFBS_ablation | 3 |
| downstream_gene_variant | 3 |
| upstream_gene_variant | 3 |
| non_coding_transcript_variant | 3 |
| NMD_transcript_variant | 3 |
| intron_variant | 3 |
| non_coding_transcript_exon_variant | 3 |
| 3_prime_UTR_variant | 5 |
| 5_prime_UTR_variant | 5 |
| mature_miRNA_variant | 5 |
| coding_sequence_variant | 5 |
| synonymous_variant | 5 |
| stop_retained_variant | 5 |
| incomplete_terminal_codon_variant | 5 |
| splice_region_variant | 5 |
| protein_altering_variant | 10 |
| missense_variant | 10 |
| inframe_deletion | 15 |
| inframe_insertion | 15 |
| transcript_amplification | 15 |
| start_lost | 15 |
| stop_lost | 15 |
| frameshift_variant | 20 |
| stop_gained | 20 |
| splice_donor_variant | 20 |
| splice_acceptor_variant | 20 |
| transcript_ablation | 20 |

**Supplementary Figure 1.**
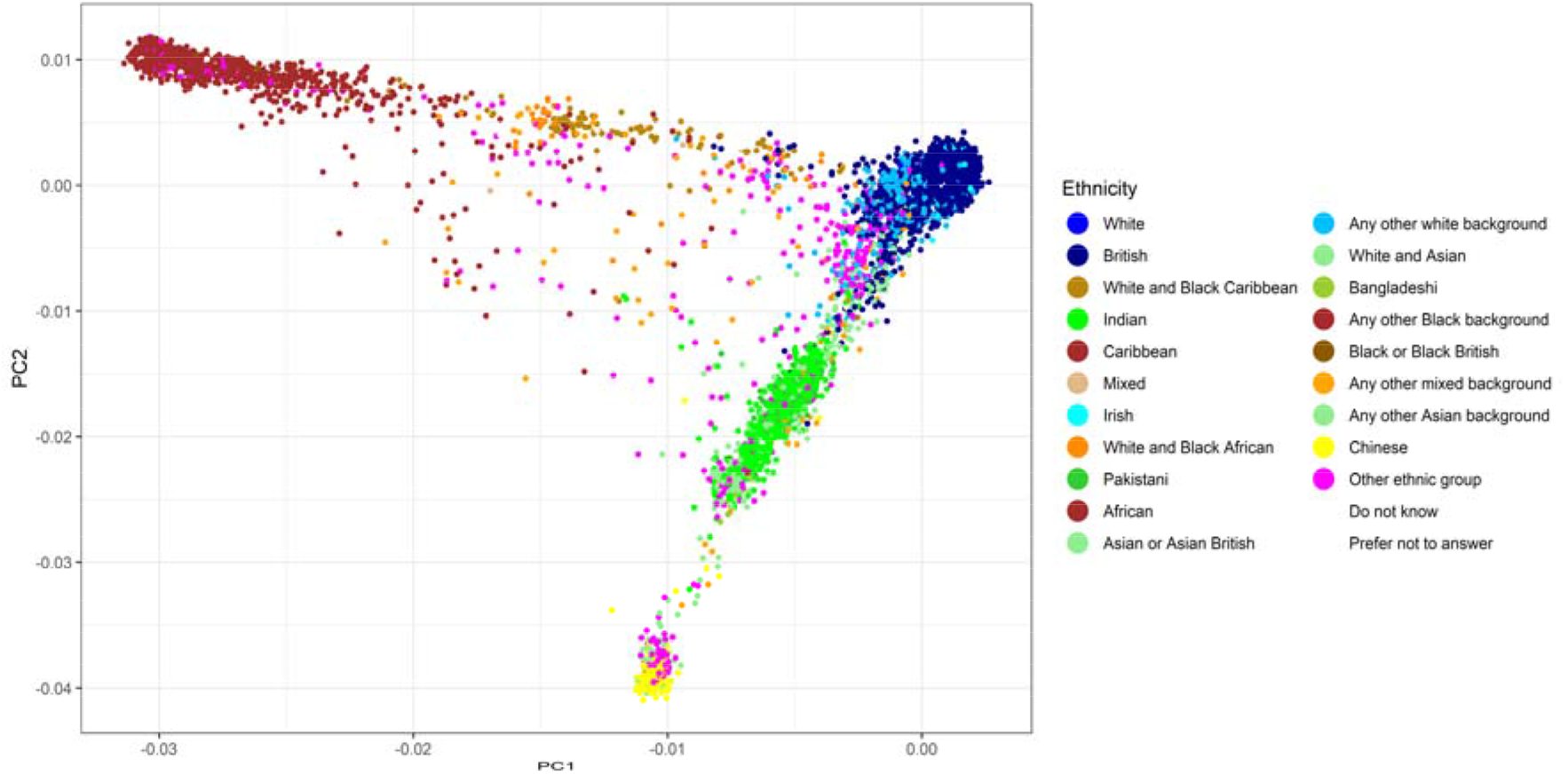
Plot of first and second principal components against self-declared ancestry.

**Supplementary Figure 2.**
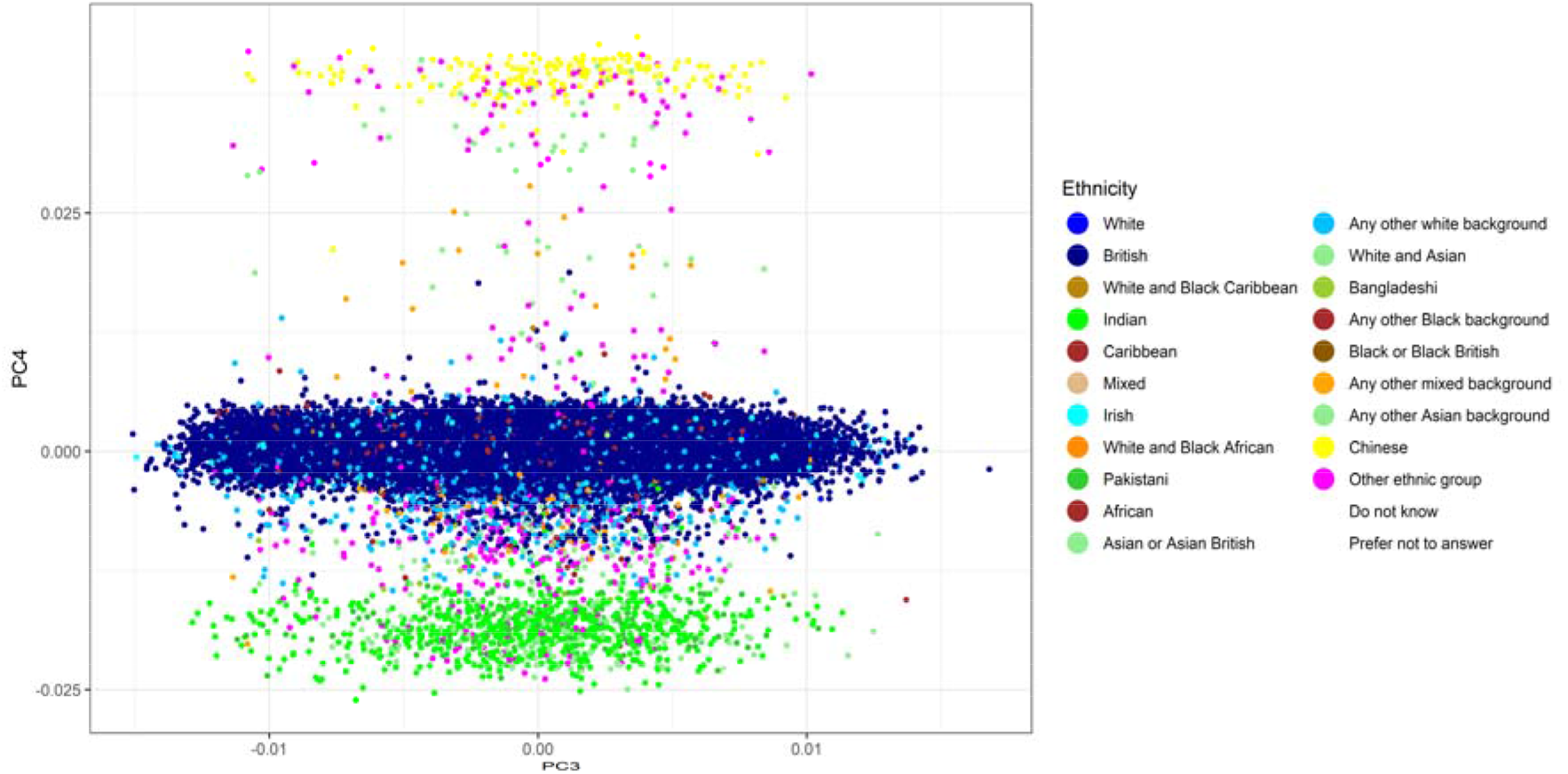
Plot of third and fourth principal components against self-declared ancestry.

**Supplementary Figure 3.**
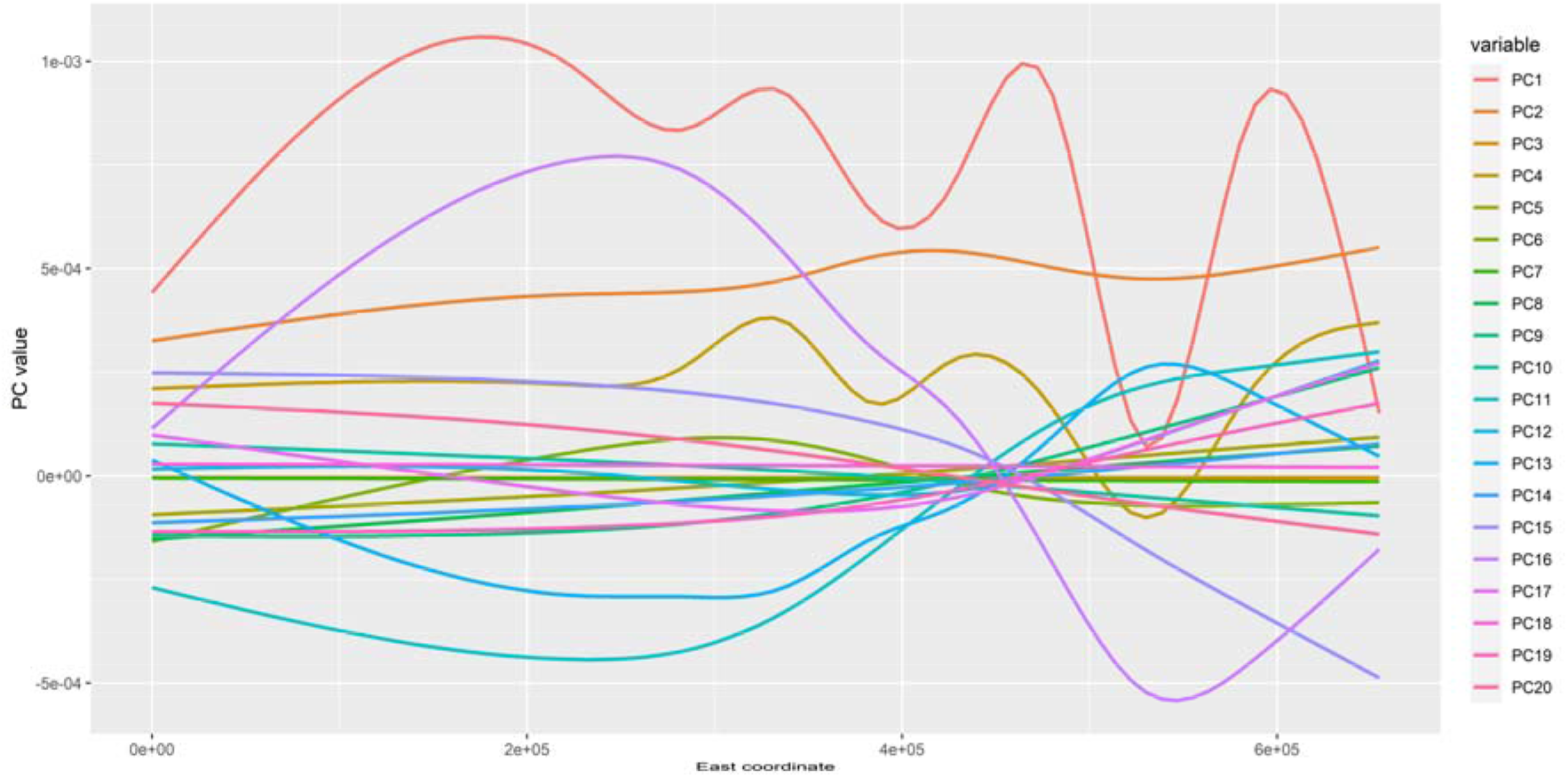
Plot of principal components against east coordinate of place of birth.

**Supplementary Figure 4.**
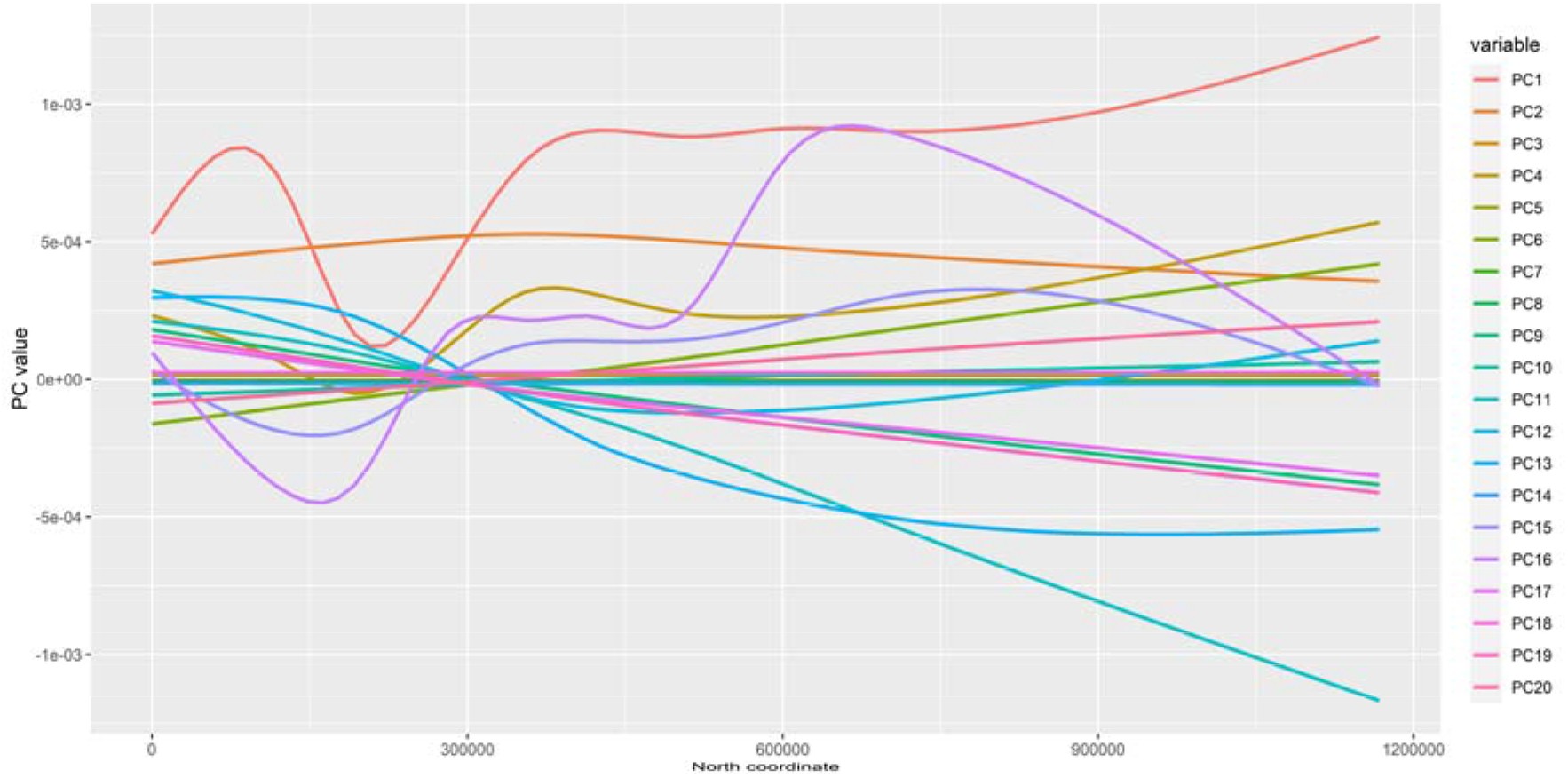
Plot of principal components against north coordinate of place of birth.

**Supplementary Figure 5.**
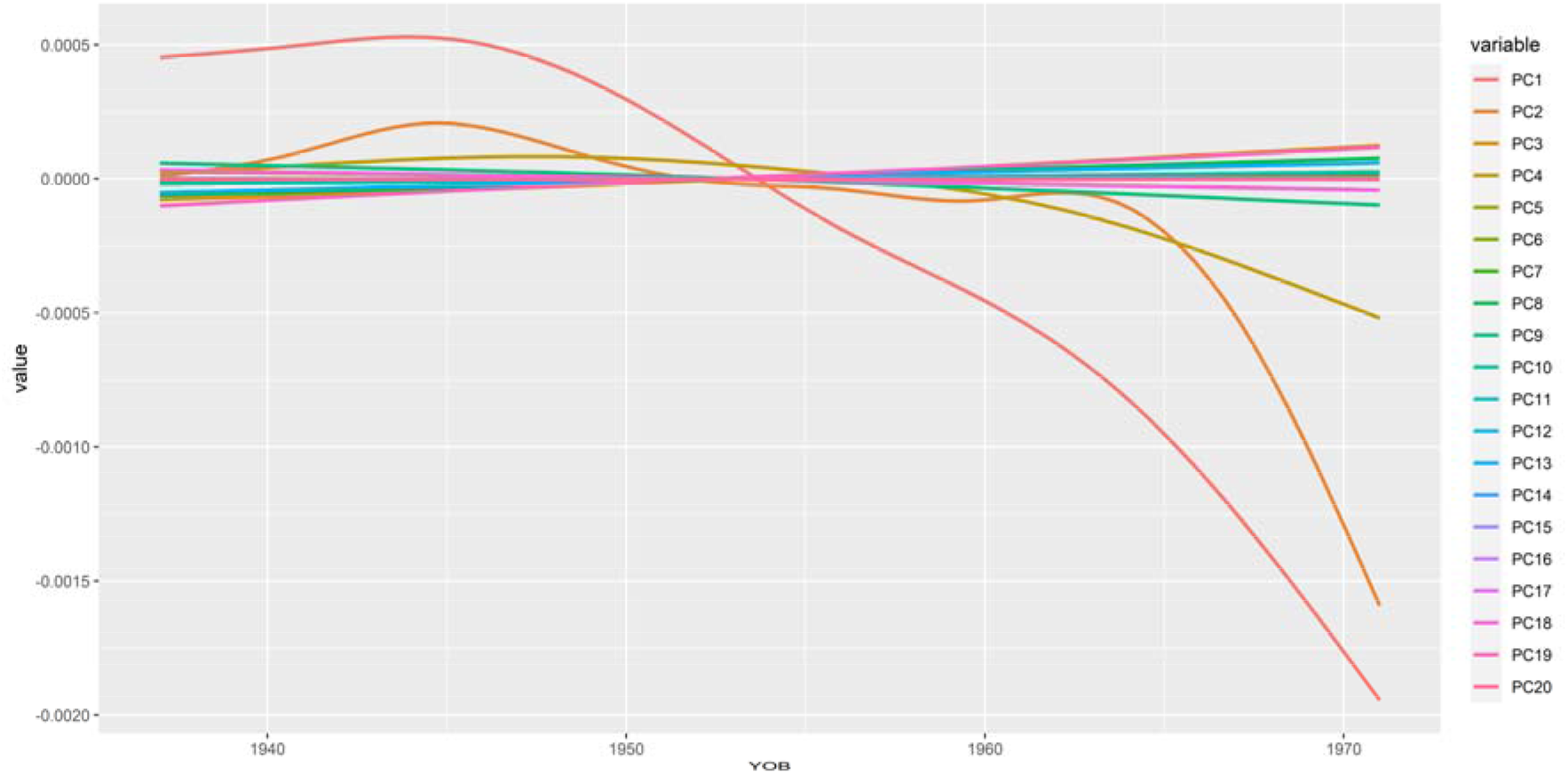
Plot of principal components against year of birth.

**Supplementary Figure 6.**
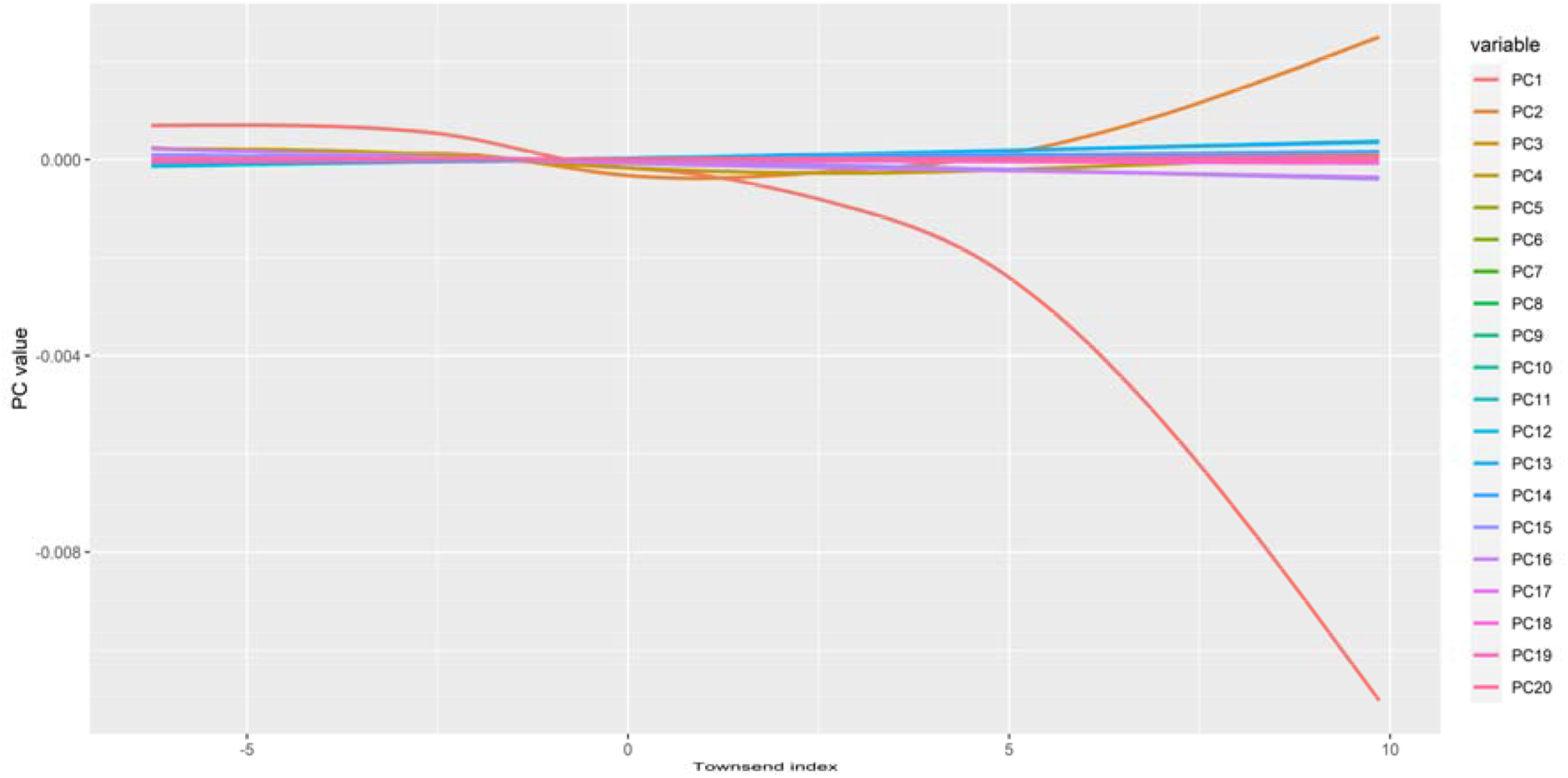
Plot of principal components against Townsend index of deprivation.

**Supplementary Figure 7.**
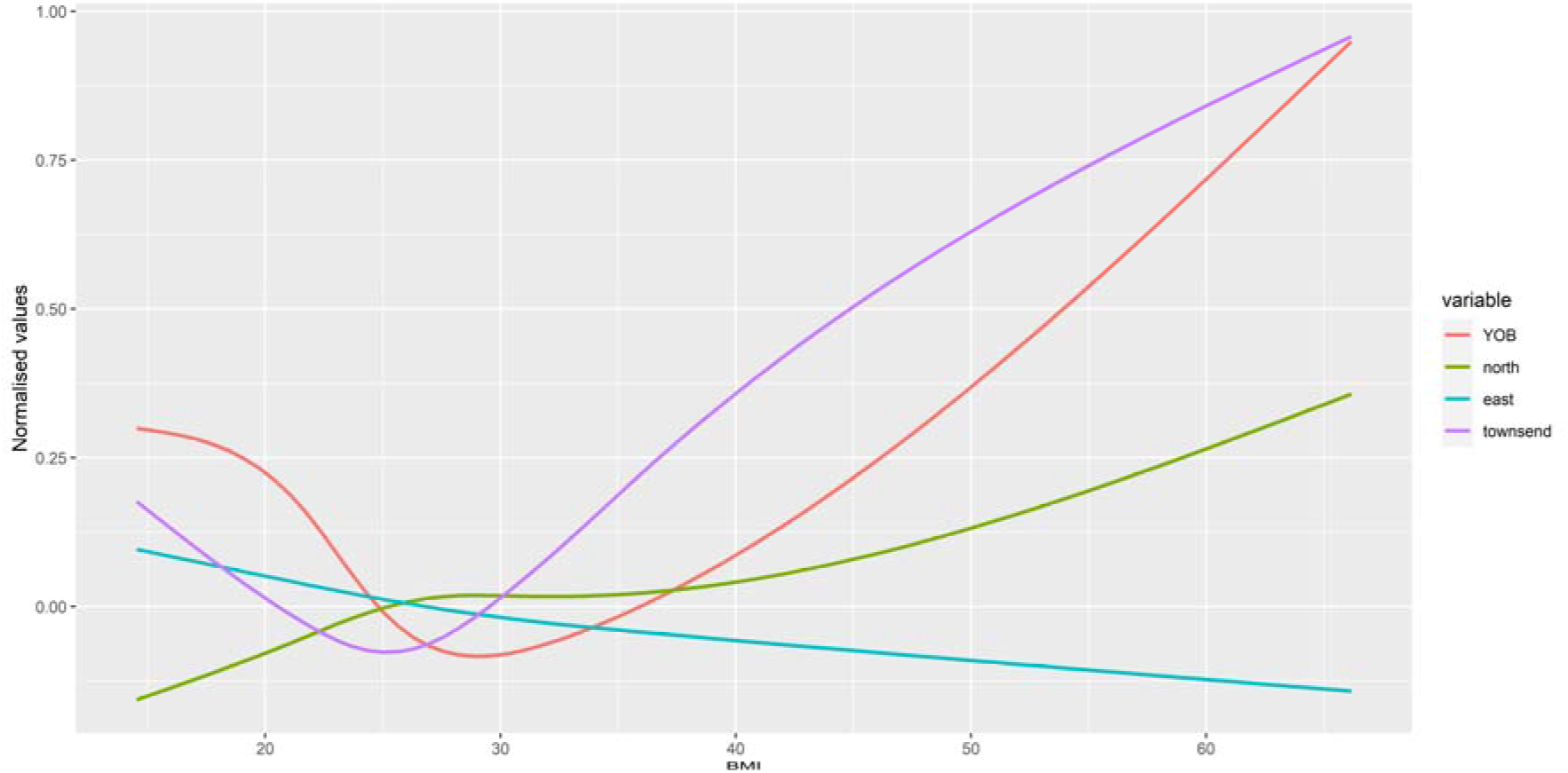
Plot of normalised values for birth coordinates, year of birth and Townsend index of deprivation against BMI.

**Supplementary Figure 8.**
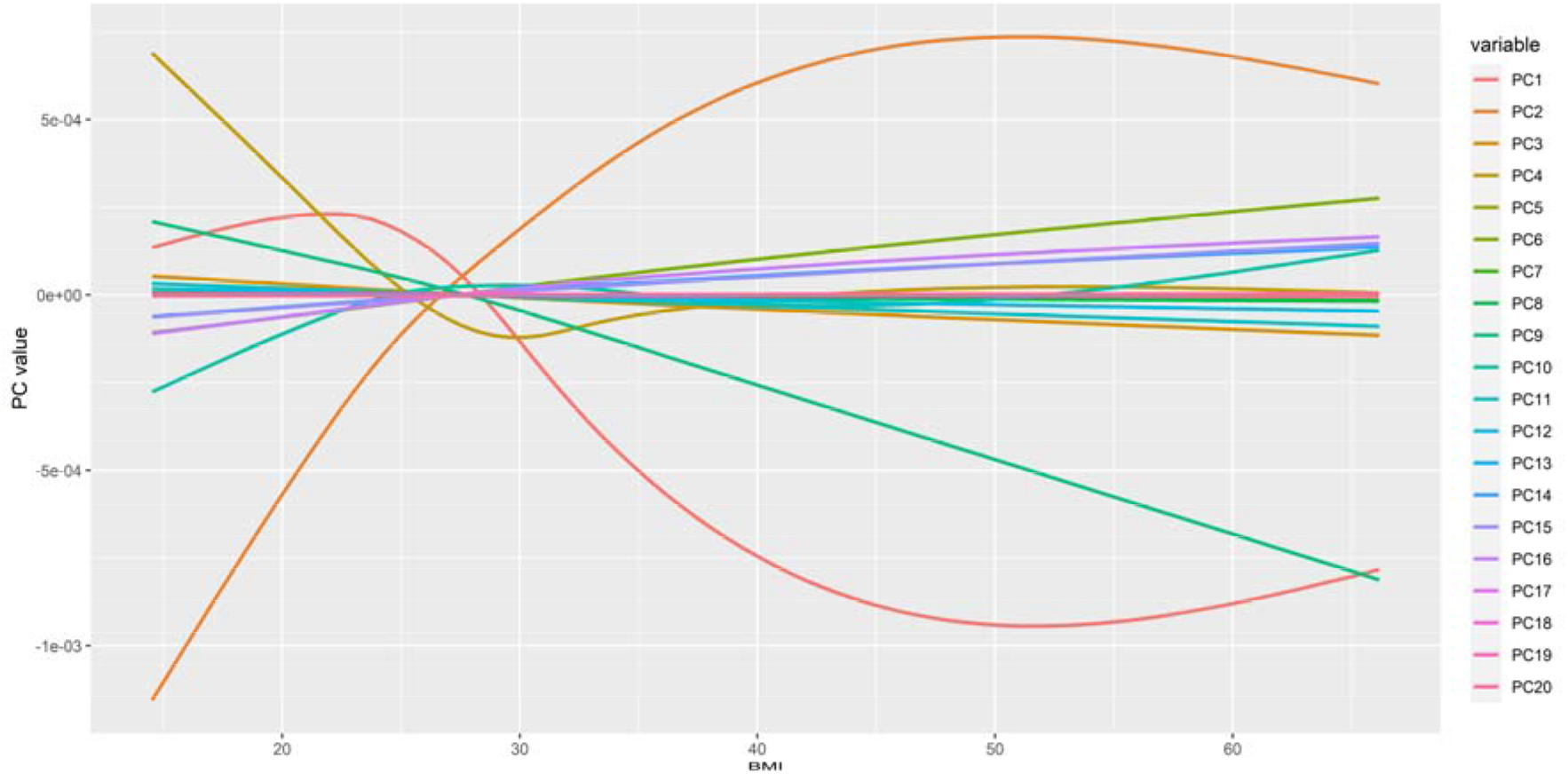
Plot of principal components against BMI.

## References

Adzhubei, I., Jordan, D.M., Sunyaev, S.R. (2013) Predicting functional effect of human missense mutations using PolyPhen-2. Curr. Protoc. Hum. Genet. 7 Unit7.20.

Bautz, D.J., Broman, K.W., Threadgill, D.W. (2013) Identification of a novel polymorphism in X-linked sterol-4-alpha-carboxylate 3-dehydrogenase (Nsdhl) associated with reduced high-density lipoprotein cholesterol levels in i/LnJ mice. G3 Genes, Genomes, Genet. 3, 1819–1825.

Bian, L., Hanson, R.L., Muller, Y.L., Ma, L., Kobes, S., Knowler, W.C., Bogardus, C., Baier, L.J. (2010) Variants in ACAD10 are associated with type 2 diabetes, insulin resistance and lipid oxidation in Pima Indians. Diabetologia 53, 1349–1353.

Bloom, K., Mohsen, A.W., Karunanidhi, A., El Demellawy, D., Reyes-Múgica, M., Wang, Y., Ghaloul-Gonzalez, L., Otsubo, C., Tobita, K., Muzumdar, R., Gong, Z., Tas, E., Basu, S., Chen, J., Bennett, M., Hoppel, C., Vockley, J. (2018) Investigating the link of ACAD10 deficiency to type 2 diabetes mellitus. J. Inherit. Metab. Dis. 41, 49–57.

Chang, C.C., Chow, C.C., Tellier, L.C., Vattikuti, S., Purcell, S.M., Lee, J.J. (2015) Second-generation PLINK: rising to the challenge of larger and richer datasets. Gigascience 4, 7.

Chen, D., Wu, P., Yang, Q., Wang, K., Zhou, J., Yang, X., Jiang, A., Shen, L., Iao, W., Jiang, Y., Zhu, L., Li, X., Tang, G. (2019) Genome-wide association study for backfat thickness at 100 kg and loin muscle thickness in domestic pigs based on genotyping by sequencing. Physiol. Genomics 51, 261–266.

Chen, Z., Yu, H., Shi, X., Warren, C.R., Lotta, L.A., Friesen, M., Meissner, T.B., Langenberg, C., Wabitsch, M., Wareham, N., Benson, M.D., Gerszten, R.E., Cowan, C.A. (2020) Functional Screening of Candidate Causal Genes for Insulin Resistance in Human Preadipocytes and Adipocytes. Circ. Res. 126, 330–346.

Cirulli, E.T., White, S., Read, R.W., Elhanan, G., Metcalf, W.J., Tanudjaja, F., Fath, D.M., Sandoval, E., Isaksson, M., Schlauch, K.A., Grzymski, J.J., Lu, J.T., Washington, N.L. (2020) Genome-wide rare variant analysis for thousands of phenotypes in over 70,000 exomes from two cohorts. Nat. Commun. 11, 1–10.

Curtis, D. (2012) A rapid method for combined analysis of common and rare variants at the level of a region, gene, or pathway. Adv Appl Bioinform Chem 5, 1–9.

Curtis, D. (2016) Pathway analysis of whole exome sequence data provides further support for the involvement of histone modification in the aetiology of schizophrenia. Psychiatr. Genet. 26, 223–7.

Curtis, D. (2019) A weighted burden test using logistic regression for integrated analysis of sequence variants, copy number variants and polygenic risk score. Eur. J. Hum. Genet. 27, 114–124.

Curtis, D., Bakaya, K., Sharma, L, Bandyopadhay, S. (2019) Weighted burden analysis of exome-sequenced late onset Alzheimer’s cases and controls provides further evidence for involvement of PSEN1 and demonstrates protective role for variants in tyrosine phosphatase genes. Ann Hum Genet 84, 291–302.

Curtis, D., Balloux, F. (2020) Editorial: Topical ethical issues in the publication of human genetics research. Ann. Hum. Genet.

Curtis, D., Coelewij, L., Liu, S.-H., Humphrey, J., Mott, R. (2018) Weighted Burden Analysis of Exome-Sequenced Case-Control Sample Implicates Synaptic Genes in Schizophrenia Aetiology. Behav. Genet. 43, 198–208.

Davey, J.R., Humphrey, S.J., Junutula, J.R., Mishra, A.K., Lambright, D.G., James, D.E., Stöckli, J. (2012) TBC1D13 is a RAB35 Specific GAP that Plays an Important Role in GLUT4 Trafficking in Adipocytes. Traffic 13, 1429–1441.

Fry, A., Littlejohns, T.J., Sudlow, C., Doherty, N., Adamska, L., Sprosen, T., Collins, R., Allen, N.E. (2017) Comparison of Sociodemographic and Health-Related Characteristics of UK Biobank Participants With Those of the General Population. Am. J. Epidemiol. 186, 1026–1034.

Genovese, G., Fromer, M., Stahl, E.A., Ruderfer, D.M., Chambert, K., Landén, M., Moran, J.L., Purcell, S.M., Sklar, P., Sullivan, P.F., Hultman, C.M., McCarroll, S.A. (2016) Increased burden of ultra-rare protein-altering variants among 4,877 individuals with schizophrenia. Nat. Neurosci. 19, 1433–1441.

Hout, C.V. Van, Tachmazidou, I., Backman, J.D., Hoffman, J.X., Ye, B., Pandey, A.K., Gonzaga-Jauregui, C., Khalid, S., Liu, D., Banerjee, N., Li, A.H., Colm, O., Marcketta, A., Staples, J., Schurmann, C., Hawes, A., Maxwell, E., Barnard, L., Lopez, A., Penn, J., Habegger, L., Blumenfeld, A.L., Yadav, A., Praveen, K., Jones, M., Salerno, W.J., Chung, W.K., Surakka, I., Willer, C.J., Hveem, K., Leader, J.B., Carey, D.J., Ledbetter, D.H., Collaboration, G.-R.D., Cardon, L., Yancopoulos, G.D., Economides, A., Coppola, G., Shuldiner, A.R., Balasubramanian, S., Cantor, M., Nelson, M.R., Whittaker, J., Reid, J.G., Marchini, J., Overton, J.D., Scott, R.A., Abecasis, G., Yerges-Armstrong, L., Baras, A., Center, on behalf of the R.G. (2019) Whole exome sequencing and characterization of coding variation in 49,960 individuals in the UK Biobank. bioRxiv 572347.

Kulkarni, S.S., Karlsson, H.K.R., Szekeres, F., Chibalin, A. V., Krook, A., Zierath, J.R. (2011) Suppression of 5□-nucleotidase enzymes promotes AMP-activated protein kinase (AMPK) phosphorylation and metabolism in human and mouse skeletal muscle. J. Biol. Chem. 286, 34567–34574.

Kumar, P., Henikoff, S., Ng, P.C. (2009) Predicting the effects of coding non-synonymous variants on protein function using the SIFT algorithm. Nat. Protoc. 4, 1073–1081.

Lawson, D.J., Davies, N.M., Haworth, S., Ashraf, B., Howe, L., Crawford, A., Hemani, G., Davey Smith, G., Timpson, N.J. (2020) Is population structure in the genetic biobank era irrelevant, a challenge, or an opportunity? Hum. Genet.

Long, T., Hicks, M., Yu, H.C., Biggs, W.H., Kirkness, E.F., Menni, C., Zierer, J., Small, K.S., Mangino, M., Messier, H., Brewerton, S., Turpaz, Y., Perkins, B.A., Evans, A.M., Miller, L.A.D., Guo, L., Caskey, C.T., Schork, N.J., Garner, C., Spector, T.D., Venter, J.C., Telenti, A. (2017) Whole-genome sequencing identifies common-to-rare variants associated with human blood metabolites. Nat. Genet. 49, 568–578.

Lotta, L.A., Gulati, P., Day, F.R., Payne, F., Ongen, H., van de Bunt, M., Gaulton, K.J., Eicher, J.D., Sharp, S.J., Luan, J., De Lucia Rolfe, E., Stewart, I.D., Wheeler, E., Willems, S.M., Adams, C., Yaghootkar, H., Forouhi, N.G., Khaw, K.-T., Johnson, A.D., Semple, R.K., Frayling, T., Perry, J.R.B., Dermitzakis, E., McCarthy, M.I., Barroso, I., Wareham, N.J., Savage, D.B., Langenberg, C., O’Rahilly, S., Scott, R.A. (2017) Integrative genomic analysis implicates limited peripheral adipose storage capacity in the pathogenesis of human insulin resistance. Nat. Genet. 49, 17–26.

McLaren, W., Gil, L., Hunt, S.E., Riat, H.S., Ritchie, G.R.S., Thormann, A., Flicek, P., Cunningham, F. (2016) The Ensembl Variant Effect Predictor. Genome Biol. 17, 122.

McLarren, K.W., Severson, T.M., Du Souich, C., Stockton, D.W., Kratz, L.E., Cunningham, D., Hendson, G., Morin, R.D., Wu, D., Paul, J.E., An, J., Nelson, T.N., Chou, A., Debarber, A.E., Merkens, L.S., Michaud, J.L., Waters, P.J., Yin, J., McGillivray, B., Demos, M., Rouleau, G.A., Grzeschik, K.H., Smith, R., Tarpey, P.S., Shears, D., Schwartz, C.E., Gecz, J., Stratton, M.R., Arbour, L., Hurlburt, J., Van Allen, M.I., Herman, G.E., Zhao, Y., Moore, R., Kelley, R.I., Jones, S.J.M., Steiner, R.D., Raymond, F.L., Marra, M.A., Boerkoel, C.F. (2010) Hypomorphic temperature-sensitive alleles of NSDHL cause CK syndrome. Am. J. Hum. Genet. 87, 905–914.

Nie, J., Han, X., Shi, Y. (2013) SAD-A and AMPK kinases: The “yin and yang” regulators of mTORC1 signaling in pancreatic β cells. Cell Cycle 12, 3366–3369.

Purcell, S., Neale, B., Todd-Brown, K., Thomas, L., Ferreira, M.A.R., Bender, D., Maller, J., Sklar, P., de Bakker, P.I.W., Daly, M.J., Sham, P.C. (2007) PLINK: a tool set for whole-genome association and population-based linkage analyses. Am. J. Hum. Genet. 81, 559–75.

Purcell, S.M., Wray, N.R., Stone, J.L., Visscher, P.M., O’Donovan, M.C., Sullivan, P.F., Sklar, P., Purcell Leader, S.M., Ruderfer, D.M., McQuillin, A., Morris, D.W., O’Dushlaine, C.T., Corvin, A., Holmans, P. a, Macgregor, S., Gurling, H., Blackwood, D.H.R., Craddock, N.J., Gill, M., Hultman, C.M., Kirov, G.K., Lichtenstein, P., Muir, W.J., Owen, M.J., Pato, C.N., Scolnick, E.M., St Clair, D., Sklar Leader, P., Williams, N.M., Georgieva, L., Nikolov, I., Norton, N., Williams, H., Toncheva, D., Milanova, V., Thelander, E.F., Sullivan, P.F., Kenny, E., Quinn, E.M., Choudhury, K., Datta, S., Pimm, J., Thirumalai, S., Puri, V., Krasucki, R., Lawrence, J., Quested, D., Bass, N., Crombie, C., Fraser, G., Leh Kuan, S., Walker, N., McGhee, K. a, Pickard, B., Malloy, P., Maclean, A.W., Van Beck, M., Pato, M.T., Medeiros, H., Middleton, F., Carvalho, C., Morley, C., Fanous, A., Conti, D., Knowles, J. a, Paz Ferreira, C., Macedo, A., Helena Azevedo, M., Kirby, A.N., Ferreira, M. a R., Daly, M.J., Chambert, K., Kuruvilla, F., Gabriel, S.B., Ardlie, K., Moran, J.L. (2009) Common polygenic variation contributes to risk of schizophrenia and bipolar disorder. Nature 10, 8192–8192.

R Core Team (2014) R: A language and environment for statistical computing. R Foundation for Statistical Computing, Vienna, Austria., Austria.

Ramphul, K., Kota, V., Mejias, S.G. (2019) Child Syndrome.

Scott, R.A., Lagou, V., Welch, R.P., Wheeler, E., Montasser, M.E., Luan, J., Mägi, R., Strawbridge, R.J., Rehnberg, E., Gustafsson, S., Kanoni, S., Rasmussen-Torvik, L.J., Yengo, L., Lecoeur, C., Shungin, D., Sanna, S., Sidore, C., Johnson, P.C.D., Jukema, J.W., Johnson, T., Mahajan, A., Verweij, N., Thorleifsson, G., Hottenga, J.-J., Shah, S., Smith, A. V, Sennblad, B., Gieger, C., Salo, P., Perola, M., Timpson, N.J., Evans, D.M., Pourcain, B.S., Wu, Y., Andrews, J.S., Hui, J., Bielak, L.F., Zhao, W., Horikoshi, M., Navarro, P., Isaacs, A., O’Connell, J.R., Stirrups, K., Vitart, V., Hayward, C., Esko, T., Mihailov, E., Fraser, R.M., Fall, T., Voight, B.F., Raychaudhuri, S., Chen, H., Lindgren, C.M., Morris, A.P., Rayner, N.W., Robertson, N., Rybin, D., Liu, C.-T., Beckmann, J.S., Willems, S.M., Chines, P.S., Jackson, A.U., Kang, H.M., Stringham, H.M., Song, K., Tanaka, T., Peden, J.F., Goel, A., Hicks, A.A., An, P., Müller-Nurasyid, M., Franco-Cereceda, A., Folkersen, L., Marullo, L., Jansen, H., Oldehinkel, A.J., Bruinenberg, M., Pankow, J.S., North, K.E., Forouhi, N.G., Loos, R.J.F., Edkins, S., Varga, T. V, Hallmans, G., Oksa, H., Antonella, M., Nagaraja, R., Trompet, S., Ford, I., Bakker, S.J.L., Kong, A., Kumari, M., Gigante, B., Herder, C., Munroe, P.B., Caulfield, M., Antti, J., Mangino, M., Small, K., Miljkovic, I., Liu, Y., Atalay, M., Kiess, W., James, A.L., Rivadeneira, F., Uitterlinden, A.G., Palmer, C.N.A., Doney, A.S.F., Willemsen, G., Smit, J.H., Campbell, S., Polasek, O., Bonnycastle, L.L., Hercberg, S., Dimitriou, M., Bolton, J.L., Fowkes, G.R., Kovacs, P., Lindström, J., Zemunik, T., Bandinelli, S., Wild, S.H., Basart, H. V, Rathmann, W., Grallert, H., Maerz, W., Kleber, M.E., Boehm, B.O., Peters, A., Pramstaller, P.P., Province, M.A., Borecki, I.B., Hastie, N.D., Rudan, I., Campbell, H., Watkins, H., Farrall, M., Stumvoll, M., Ferrucci, L., Waterworth, D.M., Bergman, R.N., Collins, F.S., Tuomilehto, J., Watanabe, R.M., de Geus, E.J.C., Penninx, B.W., Hofman, A., Oostra, B.A., Psaty, B.M., Vollenweider, P., Wilson, J.F., Wright, A.F., Hovingh, G.K., Metspalu, A., Uusitupa, M., Magnusson, P.K.E., Kyvik, K.O., Kaprio, J., Price, J.F., Dedoussis, G. V, Deloukas, P., Meneton, P., Lind, L., Boehnke, M., Shuldiner, A.R., van Duijn, C.M., Morris, A.D., Toenjes, A., Peyser, P.A., Beilby, J.P., Körner, A., Kuusisto, J., Laakso, M., Bornstein, S.R., Schwarz, P.E.H., Lakka, T.A., Rauramaa, R., Adair, L.S., Smith, G.D., Spector, T.D., Illig, T., de Faire, U., Hamsten, A., Gudnason, V., Kivimaki, M., Hingorani, A., Keinanen-Kiukaanniemi, S.M., Saaristo, T.E., Boomsma, D.I., Stefansson, K., van der Harst, P., Dupuis, J., Pedersen, N.L., Sattar, N., Harris, T.B., Cucca, F., Ripatti, S., Salomaa, V., Mohlke, K.L., Balkau, B., Froguel, P., Pouta, A., Jarvelin, M.-R., Wareham, N.J., Bouatia-Naji, N., McCarthy, M.I., Franks, P.W., Meigs, J.B., Teslovich, T.M., Florez, J.C., Langenberg, C., Ingelsson, E., Prokopenko, I., Barroso, I. (2012) Large-scale association analyses identify new loci influencing glycemic traits and provide insight into the underlying biological pathways. Nat. Genet. 44, 991–1005.

Subramanian, A., Tamayo, P., Mootha, V.K., Mukherjee, S., Ebert, B.L., Gillette, M.A., Paulovich, A., Pomeroy, S.L., Golub, T.R., Lander, E.S., Mesirov, J.P. (2005) Gene set enrichment analysis: a knowledge-based approach for interpreting genome-wide expression profiles. Proc Natl Acad Sci U S A 102, 15545–15550.

Watson, R.A., Gates, A.S., Wynn, E.H., Calvert, F.E., Girousse, A., Lelliott, C.J., Barroso, I. (2017) Lyplal1 is dispensable for normal fat deposition in mice. DMM Dis. Model. Mech. 10, 1481–1488.

Wickham, H. (2016) ggplot2: Elegant Graphics for Data Analysis. Springer-Verlag, New York.

Xu, Y., Yang, Xiao-Lin, Yang, Xiao-Long, Ren, Y.-R., Zhuang, X.-Y., Zhang, L., Zhang, X.-F. (2020) Functional Annotations of Single-Nucleotide Polymorphism (SNP)-Based and Gene-Based Genome-Wide Association Studies Show Genes Affecting Keratitis Susceptibility. Med. Sci. Monit. 26.

Zhao, Z., Bi, W., Zhou, W., VandeHaar, P., Fritsche, L.G., Lee, S. (2020) UK Biobank Whole-Exome Sequence Binary Phenome Analysis with Robust Region-Based Rare-Variant Test. Am. J. Hum. Genet. 106, 3–12.

Zhou, H., Sealock, J.M., Sanchez-Roige, S., Clarke, T.-K., Levey, D.F., Cheng, Z., Li, B., Polimanti, R., Kember, R.L., Smith, R.V., Thygesen, J.H., Morgan, M.Y., Atkinson, S.R., Thursz, M.R., Nyegaard, M., Mattheisen, M., Børglum, A.D., Johnson, E.C., Justice, A.C., Palmer, A.A., McQuillin, A., Davis, L.K., Edenberg, H.J., Agrawal, A., Kranzler, H.R., Gelernter, J. (2020) Genome-wide meta-analysis of problematic alcohol use in 435,563 individuals yields insights into biology and relationships with other traits. Nat. Neurosci.

